# Viral-mediated optogenetic stimulation of peripheral motor nerves in non-human primates

**DOI:** 10.1101/261925

**Authors:** Jordan J. Williams, Alan M. Watson, Alberto L. Vazquez, Andrew B. Schwartz

## Abstract

**Objective:** Reanimation of muscles paralyzed by disease states such as spinal cord injury remains a much sought after therapeutic goal of neuroprosthetic research. Optogenetic stimulation of peripheral motor nerves expressing light-sensitive opsins is a promising approach to muscle reanimation that may overcome several drawbacks of traditional methods such as functional electrical stimulation (FES). However, the utility of these methods has only been demonstrated in rodents to date, while translation to clinical practice will likely first require demonstration and refinement of these gene therapy techniques in non-human primates.

**Approach:** Three rhesus macaques were injected intramuscularly with either one or both of two optogenetic constructs (AAV6-hSyn-ChR2-eYFP and/or AAV6-hSyn-Chronos-eYFP) to transduce opsin expression in the corresponding nerves. Neuromuscular junctions were targeted for virus delivery using an electrical stimulating injection technique. Functional opsin expression was periodically evaluated up to 13 weeks post-injection by optically stimulating targeted nerves with a 472 nm fiber-coupled laser while recording electromyographic (EMG) responses.

**Main Results:** One monkey demonstrated functional expression of ChR2 at 8 weeks post-injection in each of two injected muscles, while the second monkey briefly exhibited contractions coupled to optical stimulation in a muscle injected with the Chronos construct at 10 weeks. A third monkey injected only in one muscle with the ChR2 construct showed strong optically coupled contractions at 5 ½ weeks which then disappeared by 9 weeks. EMG responses to optical stimulation of ChR2-transduced nerves demonstrated graded recruitment relative to both stimulus pulse-width and light intensity, and were able to track stimulus trains up to 16 Hz. In addition, the EMG response to prolonged stimulation showed delayed fatigue over several minutes.

**Significance:** These results demonstrate the feasibility of viral transduction of peripheral motor nerves for functional optical stimulation of motor activity in non-human primates, a variable timeline of opsin expression in a primate model closer to humans, and fundamental EMG response characteristics to optical nerve stimulation. Subsequently, they represent an important step in translating these optogenetic techniques as a clinically viable gene therapy.

## 1 Introduction

Disease states such as severe spinal cord injuries (SCIs) are often accompanied by muscle paralysis and loss of motor function. In these cases, restoration of native muscle and motor function is the ultimate goal of therapeutic interventions. Brain-Machine Interfaces (BMIs), which attempt to reroute control signals from an intact brain to a motor effector and effectively bypass the site of injury, have made great strides toward achieving this goal over the last several decades. BMIs have progressed from simple control of a computer cursor (Revechkis et al., 2015; Serruya et al., 2002; Taylor et al., 2002) to high-dimensional control of a robotic arm (Collinger et al., 2013; Wodlinger et al., 2015). While these systems provide an excellent intermediate step and can restore a significant degree of independence to patients that they may not have experienced for many years, they do not address the desired goal of native limb reanimation.

The traditional approach to reanimating paralyzed limbs is to electrically stimulate muscles or their nerves. This approach, often referred to as Functional Electrical Stimulation (FES), was previously coupled to residual movements or muscle activity to control the electrical stimulation of paralyzed muscles (Kilgore et al., 2008; Peckham et al., 2002). More recently, intramuscular FES has been coupled with BMI control signals by several groups to produce brain-controlled modulation of native muscle activity and limb movements in both non-human primates (NHPs) (Moritz et al., 2008; Ethier et al., 2012) and human subjects (Ajiboye et al., 2017). While these studies demonstrate restoration of volitional control over previously paralyzed muscles and can convey a significant increase in independence to a patient, the control offered by these systems is far from ideal naturalistic control. For example, in the study by Ajiboye et al. (Ajiboye et al., 2017), a C4-level SCI patient implanted with intracortical microelectrode arrays regained some volitional control over paralyzed arm muscles after pairing FES of those muscles with intracortical control signals. This scheme allowed the subject to reclaim certain daily functions such as feeding himself, but these movements were quite slow, taking several tens of seconds for tasks that most people would complete in a second or two. FES-mediated movements in this study also required additional hardware to support the arm against gravity. These studies demonstrate that the current state-of-the art for muscle stimulation necessitates major technological advances to approach practical relevance or even approach the performance level of other BMI-driven effectors such as robotic arms.

The difficulties highlighted by this study may be due to several inherent drawbacks of FES. For example, electrical stimulation of muscle activity often leads to non-physiological recruitment of muscle fibers. Under normal physiological activation, small, fatigue-resistant muscle fibers are recruited first at low activation levels followed by increasing recruitment of larger, fatigue-prone fibers at higher activation levels. In addition, the muscle fibers are often recruited in a random or reverse order (large fibers first followed by small) compared to physiological activation (small to large) (Henneman, 1957; Fang and Mortimer, 1991; Singh et al., 2000; Lertmanorat and Durand, 2004; Gregory and Bickel, 2005; Bickel et al., 2011). This reverse recruitment order can result in a poorly graded, sigmoidal recruitment curve in which gradually increasing electrical stimulation current initially results in a minimal increase in muscle force. However, upon reaching threshold, muscle force quickly increases in step-like fashion as recruited large fibers produce large force values, followed by a gradual increase in force as small fibers are additionally recruited with increasing current. This nearly “on-off” recruitment can make controlled stimulation of intermediate force values difficult to achieve. The reverse recruitment order elicited by FES can also lead to early fatigue of muscle contractions (Gregory and Bickel, 2005). Small diameter, fatigue-resistant motor units are normally recruited first for prolonged or repetitive but low-amplitude force production while large, fatigue-susceptible units are activated later and more sparsely for bursts of activity requiring higher force. When this relationship is reversed and large fibers are activated early and often, prolonged electrical stimulation causes the large fibers to fatigue early and the muscle is less able to produce forceful contractions. These factors have limited the use of FES for reanimation of paralyzed muscle.

A potential alternative to FES that may circumvent some of these shortcomings is the use of peripheral optogenetic techniques to elicit muscle activity through Functional Optical Stimulation (FOS). In this approach, light sensitive ion channels, i.e. “opsins”, are inserted into the motor nerve axonal membrane, allowing the nerve to be depolarized using light stimulation. FOS experiments in rodents have suggested that this approach may hold several advantages over FES by overcoming the drawbacks of FES discussed above. In an initial study, Llewellyn et al. demonstrated in transgenic mice expressing the blue-light sensitive channelrhodopsin (ChR2) that optically stimulating motor nerves elicits a natural recruitment of muscle fibers; small diameter muscle fibers are recruited first with low-amplitude stimulation, and larger fibers are recruited with increasing stimulation intensities (Llewellyn et al., 2010). This combination of fiber activation produced a wide dynamic range of forces from fine to gross. This differs from the recruitment order of muscle fibers observed with FES. Related to these observations of natural recruitment order, Llewellyn *et al.* also found that muscle activation leads to decreased muscle fatigue, delaying the onset of muscle fatigue to repetitive optical stimulation for several minutes versus only a few tens of seconds with electrical stimulation (Llewellyn et al., 2010). Additionally, viral transduction of opsins sensitive to different wavelengths of light makes it possible to selectively target only nerve fibers innervating a desired muscle. Conversely, FES is relatively non-selective in stimulating axons to individual muscles at proximal nerve sites. For example, electrically stimulating the sciatic nerve will activate contractions of the gastrocnemius and tibialis anterior muscles (as well as others) non-specifically. However, Towne *et al.* have demonstrated selective activation of viral-targeted tibialis anterior muscle with optical stimulation at a common proximal sciatic nerve location (Towne et al., 2013). Although selective electrical stimulation can be accomplished to a degree, it typically requires complex spatial activation patterns or deforming the nerve (Tyler and Durand, 2002). Finally, although optical stimulation can cause photoelectric artifacts if shone directly on an electrode (Kozai and Vazquez, 2015), it does not cause EMG artifacts to arise from distant optical stimulation, unlike the volume-conducted artifact associated with electrical stimulation of a muscle or nerve. This artifact-free stimulation could simplify signal processing in closed-loop stimulation schemes that rely on EMG or intraneural feedback (Yeom and Chang, 2010; Bruns et al., 2013). Overall, these potential advantages may make peripheral optogenetic stimulation a viable alternative to FES for muscle activation in neuroprosthetic applications.

To date, peripheral optogenetic activation of muscle activity displaying these potential benefits has only been demonstrated in rodents. Across these studies, several methods have been used to label motor nerves with stimulating opsins. Llewellyn *et al.* used a transgenic mouse line to express ChR2 in neurons in the peripheral nervous systems (PNS) under the Thy1 promoter, allowing the authors to elicit muscle activity through optical stimulation of peripheral motor nerve axons (Llewellyn et al., 2010). While use of transgenic mouse lines of this nature is useful for testing the neurophysiological characteristics of optogenetic stimulation, this type of germ-line manipulation is impractical for human applications. Conversely, Bryson *et al.* demonstrated optical control of muscle in wild-type mice after transplanting motor neurons expressing ChR2 derived from embryonic stem cells into a nerve graft site (Bryson et al., 2014). A more common approach in line with genetic manipulation used in other systems and disease models (Asokan et al., 2012) is the utilization of adeno-associated virus (AAV) vectors to enable optical modulation of muscle activity. These include expression of ChR2 in rat peripheral motor nerves following muscle injection of an AAV vector (Maimon et al., 2017; Towne et al., 2013) or expression directly in mouse skeletal muscle tissue following systemic injection (Bruegmann et al., 2015). While direct optical modulation of muscle tissue is feasible, it would require individual light sources for each targeted muscle, and implanting optical stimulation hardware could be difficult in smaller or deep muscles such as intrinsic hand muscles. Conversely, viral transduction of opsins in motor nerve axons offers the potential to independently control multiple muscles from a single proximal nerve location more amenable to light source implantation, making it an appealing approach over muscle transduction.

While the above rodent studies are an important step in exploring the potential of peripheral optogenetic stimulation, translating these techniques to NHPs prior to human trials remains largely unexplored. Indeed, even the development of optogenetic techniques for NHPs in the central nervous system (CNS) has proven challenging with examples of successful viral transduction studies slowly beginning to accumulate (Cavanaugh et al., 2012; Dai et al., 2014; Diester et al., 2011; El-Shamayleh et al., 2017; Han et al., 2009; Jazayeri et al., 2012; Stauffer et al., 2016). A handful of studies have demonstrated viral transduction of peripheral neuromuscular tissue with fluorescent proteins in NHP models (Okada et al., 2013; Towne et al., 2009), but optogenetic modulation of peripheral neural activity in primates similar to the aforementioned rodent studies has not been reported to date. With the difficulties in translating rodent-proven optogenetic techniques to primates, it is likely that similar challenges will be faced in translating peripheral motor optogenetic techniques due to the PNS’s greater exposure to the immune system, the differences in rodent vs. primate immune responses, and sheer scale difference.

Given the potential benefits of FOS over current FES approaches for neuroprosthetic applications and the current gap regarding peripheral optogenetic modulation of motor activity in higher-order animal models, the current study examined the feasibility of virally mediated optogenetic modulation of motor activity in a macaque model. We hypothesized that we could utilize an AAV vector with prior success in rodent FOS studies to transduce macaque motor nerves with commonly used opsins (ChR2 and Chronos) and drive PNS motor activity in an NHP model. To overcome some of the aforementioned translational challenges, we delivered the virus using a stimulating muscle injection technique to target and deliver virus locally near neuromuscular junctions. We then used optical stimulation of targeted nerves and electromyographic (EMG) recordings to functionally assess opsin expression. We next examined the relationships between optical stimulation variables and elicited EMG activity for comparison with previous observations in rodents. Finally, we examined histological and whole tissue imaging of opsin expression to correlate expression variability with observations of functional optical sensitivity in nerve samples. The results presented will not only help to address the feasibility of peripheral viral gene therapy and FOS in BMI applications, but they may also help to identify further virus, opsin, and hardware development needed prior to clinical translation.

## 2 Methods

### 2.1 Subjects

For the main focus of this study, three male rhesus macaques (Macaca mulatta), Monkeys M, O, and P, weighing 7-9 kg were used in these experiments. In a set of preliminary experiments, we injected viral constructs into male Fischer 344 rats weighing 200-400 g to serve as a baseline for comparison with our non-human primate experiments (see Supplementary Methods). All animal procedures were approved and conducted in accordance with the University of Pittsburgh’s Institutional Animal Care and Use Committee.

### 2.2 Viral constructs

Two high-titer AAV6-based viral vectors were obtained from Virovek, Inc. (Hayward, CA) for use in these experiments. The first vector, AAV6-hSyn-ChR2(H134R)-eYFP, was produced at a titer of 1.04x10^14^ vp/mL. This construct was previously used for AAV-mediated transduction of excitatory opsins in peripheral motor nerves (Maimon et al., 2017; Towne et al., 2013), and has been reported at a range of viral titers. The second construct tested in these experiments replaced the well-studied opsin, ChR2(H134R), with the more recently developed Chronos (Klapoetke et al., 2014). Chronos has faster kinetics and increased sensitivity over ChR2, but its utility has not been demonstrated in the periphery to date. The AAV6-hSyn-Chronos-eYFP construct was produced by Virovek at a titer of 1.00x10^14^ vp/mL.

### 2.3 Virus injections

Aseptic techniques were used for all virus injection surgeries. Prior to virus injection, each monkey was sedated with a cocktail of ketamine (20 mg/kg) and xylazine (0.5 mg/kg). For each target muscle, a skin incision was made to expose the muscle while leaving the surrounding fascia intact. Following virus injection procedures as described below, all skin incisions were closed with subcuticular stitches. Injected animals received a 5 day course of antibiotics and were returned to their home cage to recover for at least three weeks before evaluation expression.

Monkeys M, O, and P received injections of AAV-based constructs in two, four, and one muscle(s), respectively, as shown in Figure 1. Monkey M was injected in two muscles with the AAV6-hSyn-ChR2(H134R)-eYFP construct. The tibialis anterior (TA) muscle of each leg was injected with construct diluted to relatively low or high titer. The right TA (“high” titer leg) was injected with 160 μL of virus (1.66x10^13^ vp) diluted in hypertonic saline to a total volume of 2 mL (8.32x10^12^ vp/mL). The left TA (“low” titer) was injected with 20 μL of virus (2.08x10^12^ vp) diluted with hypertonic saline to a volume of 2 mL (1.04x10^12^ vp/mL). Ten individual injections were made per muscle with approximately 200 μL of virus solution injected per site.

**Figure 1.**
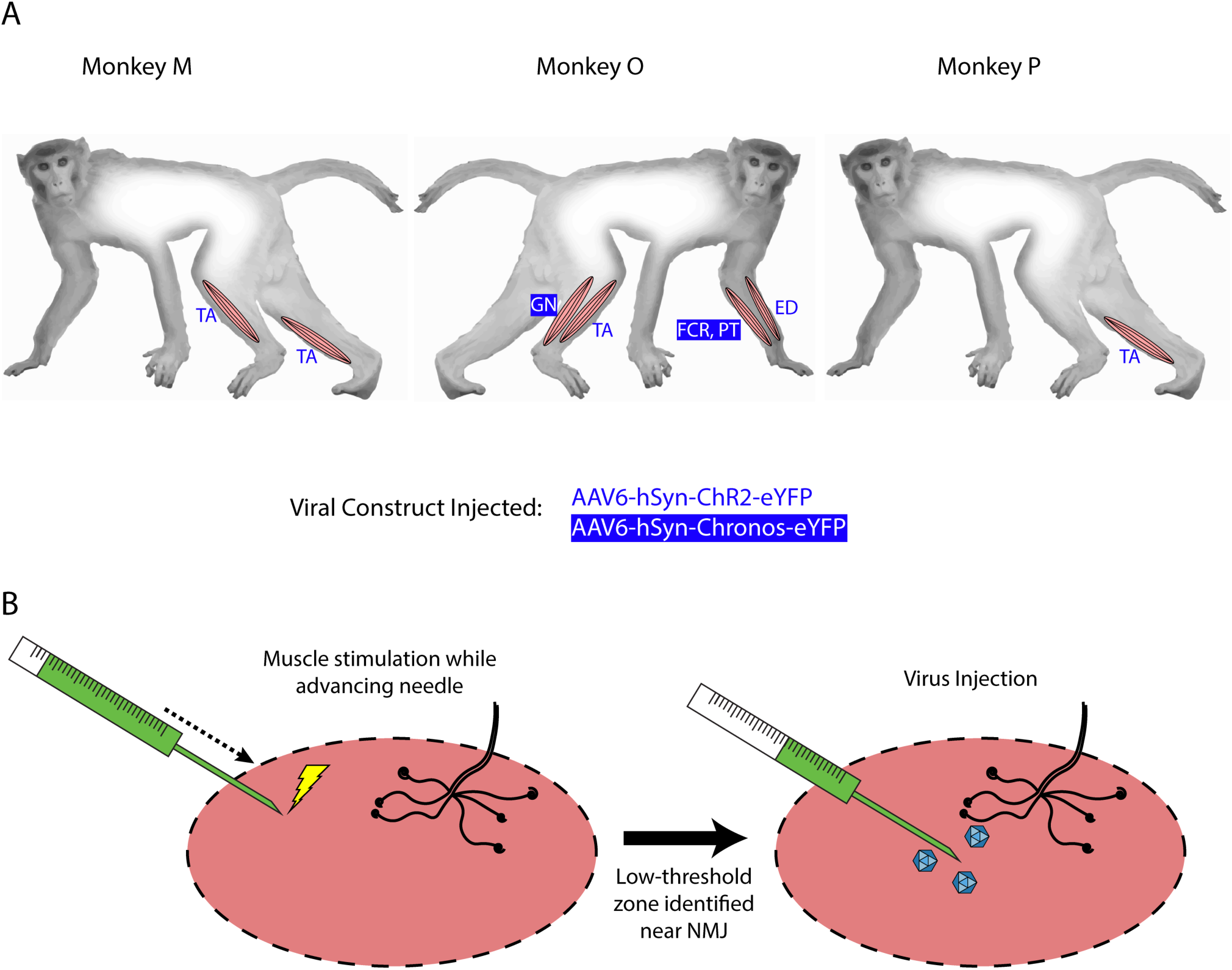
Intramuscular injections of viral optogenetic constructs. A) *Schematic of macaque subjects, viral constructs, and muscles injected*. Blue/reverse blue lettering indicate whether a designated muscle was injected with the ChR2-or Chronos-based construct, respectively. Muscle abbreviations: TA – tibialis anterior, GN – lateral gastrocnemius, FCR – flexor carpi radialis, PT – pronator teres, ED – extensor digitorum. B) *Identification of injection zones*. A stimulating injection needle was advanced slowly into the muscle while applying low-amplitude electrical stimulation. After observing near-maximal contractions, indicating a zone with higher density of neuromuscular junctions (NMJs), 200 mL of viral solution was injected slowly over 1 minute. This process was repeated over ten sites per muscle or muscle group.

At each injection site, low-threshold electrical stimulation was used to localize potential motor endplates to minimize the distance virus would have to diffuse before uptake at the neuromuscular junction. A 30 gauge monopolar injectable needle (Technomed, Netherlands) was attached to a tuberculin syringe filled with virus solution, while a metal hub needle attached to a ground lead was inserted through the skin edge. A biphasic waveform (200 μs at 0.25 mA, 400 μs at -0.125 mA) was applied between the needle tip and ground electrode via an analog stimulus isolator (A-M Systems, Model 2200). As electrical stimulation was applied, the needle was slowly advanced into the muscle by hand while monitoring muscle twitches. After finding a needle insertion position facilitating maximum contraction, stimulation was paused and 200 μL of virus were injected over approximately 1 minute. The needle was held in place for an additional minute before slowly withdrawing it. This process was repeated for each injection site. Injections were aimed at the presumed line of neuromuscular junctions approximately 1/3 of the muscle length away from the proximal end of the muscle. Needle insertions were aimed in both proximal-to-distal and distal-to-proximal fashions toward this zone, and were spaced laterally across the muscle surface.

Monkey O received injections of both the ChR2 and Chronos viral constructs. Four muscles groups (two flexor/extensor pairs) were targeted. In the right leg, we injected TA muscle with the AAV6-hSyn-ChR2-eYFP construct, and we injected the lateral gastrocnemius (GN) with AAV6-hSyn-Chronos-eYFP. In the left forearm, we injected the extensor digitorum (ED) with the ChR2 construct, and we injected both flexor carpi radialis (FCR) and pronator teres (PT) muscles with the Chronos solution. For each muscle, 100 μL of stock virus was diluted with hypertonic saline to 2 mL total volume (5.02x10^12^ vp/mL for ChR2, 5.0x10^12^ vp/mL for Chronos). The Chronos solution was split evenly between the FCR and PT muscles in the forearm. Targeting of the muscle endplates and muscle injections were performed in a similar fashion to those described for Monkey M.

Monkey P was injected with the AAV6-ChR2 construct in the right TA muscle. 100 μL of stock virus (1.04x10^13^ vp) was diluted to a total volume of 1 mL with hypertonic saline at a slightly higher concentration (1.04x10^13^ vp/mL) than the highest used used in Monkey M. Stimulating injections targeting neuromuscular junctions were used to deliver 900 μL of virus solution to the muscle over 5 sites. The deep peroneal (DP) nerve innervating the TA muscle was also exposed near its insertion into the TA via blunt separation of fibers of the overlying biceps femoris muscle. 100 μL of virus solution was injected directly into the DP nerve over 3 sites.

### 2.4 Expression evaluation

Each monkey was periodically evaluated for opsin expression over the course of 8-13 weeks. During an evaluation surgery, the monkey was anesthetized, and a previously injected muscle was re-exposed. Blunt dissection was used to separate fascia from the muscle and to expose the innervating nerve. Electrical stimulation of the nerve using a pair of bipolar hook electrodes (Cadwell Laboratories, Kennewick, WA) was used to confirm the identity of the desired nerve. Optical stimulation was delivered using a 400 μm diameter core multimode fiber (ThorLabs, Newton, NJ) connected to a 150 mW, 472 nm fiber-coupled laser (LaserGlow Technologies, Toronto, Ontario). Maximum laser output at the fiber tip was typically around 110 mW. While moving the fiber tip manually along the length of the nerve, optical stimulation trains of 15-20 ms pulses at 2.5 Hz and 100 mW were delivered to scan the nerve for areas sensitive to optical stimulation. A pair of the injectable electrode needles (same model as used for stimulation during muscle injection) was inserted into the muscle belly to measure EMG activity with a metal hub needle in the skin edge serving as electrical ground. EMG electrodes were connected to a low-impedance differential headstage with 20x gain (RA16LI-D, Tucker Davis Technologies (TDT), Alachua, FL). A TDT neurophysiology recording system (RZ-2) was used to coordinate optical stimulation waveforms with EMG recordings. All waveforms were sampled at 24 kHz.

Following periodic evaluations of nerve expression, any retracted muscle and fascia overlying target nerves were sutured in layers with absorbable suture. Skin incisions were closed with subcuticular stitches, and the animal was returned to his cage to recover.

### 2.5 Perfusion, tissue clearing, and imaging

Following final evaluation of opsin expression, each animal was perfused transcardially with 1X phosphate buffer solution (PBS) followed by 4% paraformaldehyde (PFA). Sections of targeted nerves were harvested and post-fixed in 4% PFA overnight, after which they were stored in 0.02% sodium azide solution in PBS at 4° C while awaiting processing for tissue clearing. Several nerve samples from each animal were reserved for tissue clearing and whole sample imaging. 5-10 mm long sections of nerve were excised from the main nerve branch directly innervating virus targeted muscles. Nerve samples were cleared using the polyethylene glycol (PEG)-associated solvent system (PEGASOS) passive immersion protocol (Jing et al., 2018). Briefly, following tissue fixation in 4% PFA, tissues were first passively bathed in a 25% Quadrol to decolor the tissue step, followed by gradient solutions (30%, 50%, 70%) of tert-Butanol (tB) at 37° over two days for delipidation and dehydration. Samples were further dehydrated in a solution composed of 70% tB, 27% PEG methacrylate M_n_ 500 (PEGMMA500), and 3% Quadrol for two days. Finally tissues were cleared for at least one day in a solution of BB-PEG formed by mixing 75% benzyl benzoate (BB) and 25% PEGMMA500 supplemented with 3% Quadrol until tissues reached transparency. Following clearing, samples were preserved in the BB-PEG clearing medium at room temperature.

Following tissue clearing, whole nerve samples were mounted in BB-PEG and sealed between two rounded cover glass. Tissues were imaged using the RS-G4 ribbon scanning confocal microscope (Caliber I.D., Rochester, NY) (Watson et al., 2017) equipped with an iChrome MLE laser engine (Toptica Photonics, Munich Germany). Large-area mosaic images were captured using the Olympus XLPLN25XWMP2, 25x, 1.05NA, water immersion objective with a scan zoom of 1.7 and lateral resolution of 0.295 microns. Z-steps were acquired at 1.52 microns. Fluorescence from eYFP was detected by using 488 nm excitation and 520/44 nm emission filters. The tissue was imaged from bottom to top with the 488-laser power interpolated linearly through Z from 15% (top) to 30% (bottom). Mosaic images were stitched and assembled by the microscope software.

To facilitate analysis, mosaic images were processed in MATLAB R2017b by first flattening the image and then subtracting background. A unique background filter was calculated for each image by applying a gaussian filter with a standard deviation of 6 and then a morphologic opening with a disk structuring element of radius 200 pixels. A flattening filter was produced by taking the mean of the background filter divided by the background filter:

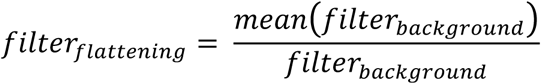

First the flattening filter was applied to both the RAW image and the background filter by multiplying the two images. The flattened background filter was subtracted from the resulting image:

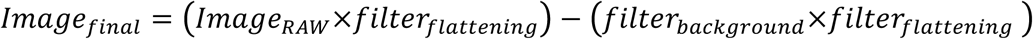

Flattened and background-subtracted images were then assembled into volumes using the Imaris File Converter and analyzed using Imaris v9.2.1 (Bitplane). Models of eYFP expressing nerve axon segments were built using the Imaris Surpass surface tool. Manual cleaning of the surface rendering was performed to ensure that labeled regions represented eYFP expressing nerves. Volume and position data for eYFP model surface elements were exported from the surface tool to Matlab where expression volume was binned as a function of longitudinal (Y) position.

## 3 Results

### 3.1 Time course of expression

Each monkey was tested intermittently for expression in targeted nerves intermittently between the injection surgery and final terminal evaluation surgery. Monkey M was tested at 3 (right TA), 4 (left TA), and 8 weeks (both legs). During evaluation time points at week 3 and 4, neither leg showed visible contractions or EMG deflections upon laser stimulation at full power. The deep peroneal (DP) nerve innervating the TA muscle was exposed for stimulation but was not aggressively dissected to avoid permanently damaging the nerve. At week 8, both targeted nerves were re-exposed and tested. Initial optical stimulation along the length of the superficial (anterior) portion of the left DP nerve again did not suggest overt expression of ChR2. However, stimulation of the posteriolateral aspect of the nerve at a single proximal site demonstrated visible contraction of the TA muscle. Optical stimulation of the anterior aspect of the nerve at this location or proximal/distal to it did not elicit muscle contractions. However, following this initial display of sensitivity, the DP was dissected distally to its insertion into the TA muscle where it branches out. At this point, optical stimulation of the nerve and muscle activity became more consistent with several branches showing sensitivity. Evaluation of the DP nerve of the right leg proceeded in a similar fashion with dispersed sensitive spots along the nerve proximal to the muscle, and more consistent sensitivity where the nerve branched out close to the muscle.

Expression of optogenetic transduction in Monkey O was tested at 5, 10, and 13 weeks post-injection. We first tested expression in the nerves leading to the TA and lateral GN muscles of the right leg at 5 weeks. No visible muscle contractions were elicited with optical stimulation of either nerve. At 10 weeks, we tested all targeted nerves in the right leg and left forearm. Neither nerve branch in the leg nor nerve supplying the ED muscle in the forearm demonstrated optically sensitivity. A branch of the median nerve supplying the PT muscle in the left forearm (injected with the Chronos vector) did facilitate brisk contraction of the PT when stimulated optically with the fiber-coupled laser. Upon observing optical sensitivity, we further dissected the nerve to accommodate placement of an LED nerve cuff intended for chronic stimulation of the nerve. After initial placement of the cuff, the nerve no longer initiated PT contractions when stimulated with blue light from the cuff or optical fiber. We suspected the nerve may have become irritated by prolonged exposure or irritation during the LED cuff placement, so we removed the cuff, re-sutured all nerve and muscle exposures, and returned the monkey to its home cage. At 13 weeks, we re-tested each targeted nerve. During this experiment, no nerves (including the previously sensitive branch to the PT muscle) exhibited optical sensitivity. Electrical stimulation of the PT muscle’s nerve elicited brisk contractions, suggesting the nerve’s health was intact.

The DP nerve of Monkey P innervating the injected right TA muscle was tested at 5 ½ weeks and 9 weeks post-injection. At the first checkpoint, optical stimulation of the exposed nerve resulted in small contractions visible through the skin. Optical sensitivity was more consistent along the exposed portion of the nerve than in Monkeys M and O with no obvious insensitive portions of nerve. After returning to check the nerve at the 9 week time point, no visual or EMG evidence of sensitivity to optical stimulation of the target nerve was present.

### 3.2 Visually observed responses to optical stimulation

Visible contractions to optical stimulation of targeted nerves were observed in all three monkeys. In monkey M, contractions of the TA muscle were clearly visible in both legs. Additionally, contractions of different portions of the muscle could be observed when different fascicles were stimulated at the branch-out location of the nerve near insertion into the muscle. However, even at full power stimulation (>100 mW, 30 ms pulse duration), optical stimulation along the nerve did not produce functional movement of the lower leg (i.e. dorsiflexion of the foot). For the short period of time that we observed optical sensitivity in monkey O, optical stimulation of a branch of the median nerve produced brisk contractions of the PT muscle similar to those observed in monkey M. Again, although clearly visible, these contractions did not result in pronation of the forearm. Finally, optical stimulation of the right DP nerve in Monkey P resulted in contractions of the TA that could be seen through the skin before further exposing the muscle belly.

As a set of visual checks that opsin expression was limited to nerve tissue innervating the target muscle, no muscle contractions were observed when the injected muscle was directly stimulated with blue light. In addition, optical stimulation of nearby non-injected muscles and their corresponding nerves did not induce visible contractions or EMG activity.

### 3.3 Individual optical pulses elicit graded EMG responses to pulse duration and intensity

After observing visual responses to optical stimulation in monkey M, we recorded the EMG response of each TA muscle to variations in several optical stimulation parameters. First, the pulse duration was varied from 1 to 30 ms (100 mW). The delay from the onset of the optical pulse to a deflection in EMG activity was consistent across muscles at approximately 12 ms. As shown in Figure 2A, the length of the evoked EMG waveform stays relatively constant while the peak amplitude and RMS increase gradually with pulse duration until plateauing at pulse durations above 10 ms as shown in Figure 2B. We then measured the evoked EMG activity as a function of the incident intensity of optical stimulation. Figure 2D depicts a near-linear monotonic increase of EMG activity with optical intensity within the range studied. Results from Monkey P showed similar trends with a plateau in elicited EMG activity near 10 ms and a linear increase in EMG with light intensity (see Supplemental Figure S1a-b). These trends are consistent with results from previous studies in rodents utilizing AAV6 and ChR2 (Llewellyn et al., 2010; Towne et al., 2013), and support the notion that optogenetic stimulation offers graded recruitment of muscle activity.

**Figure 2.**
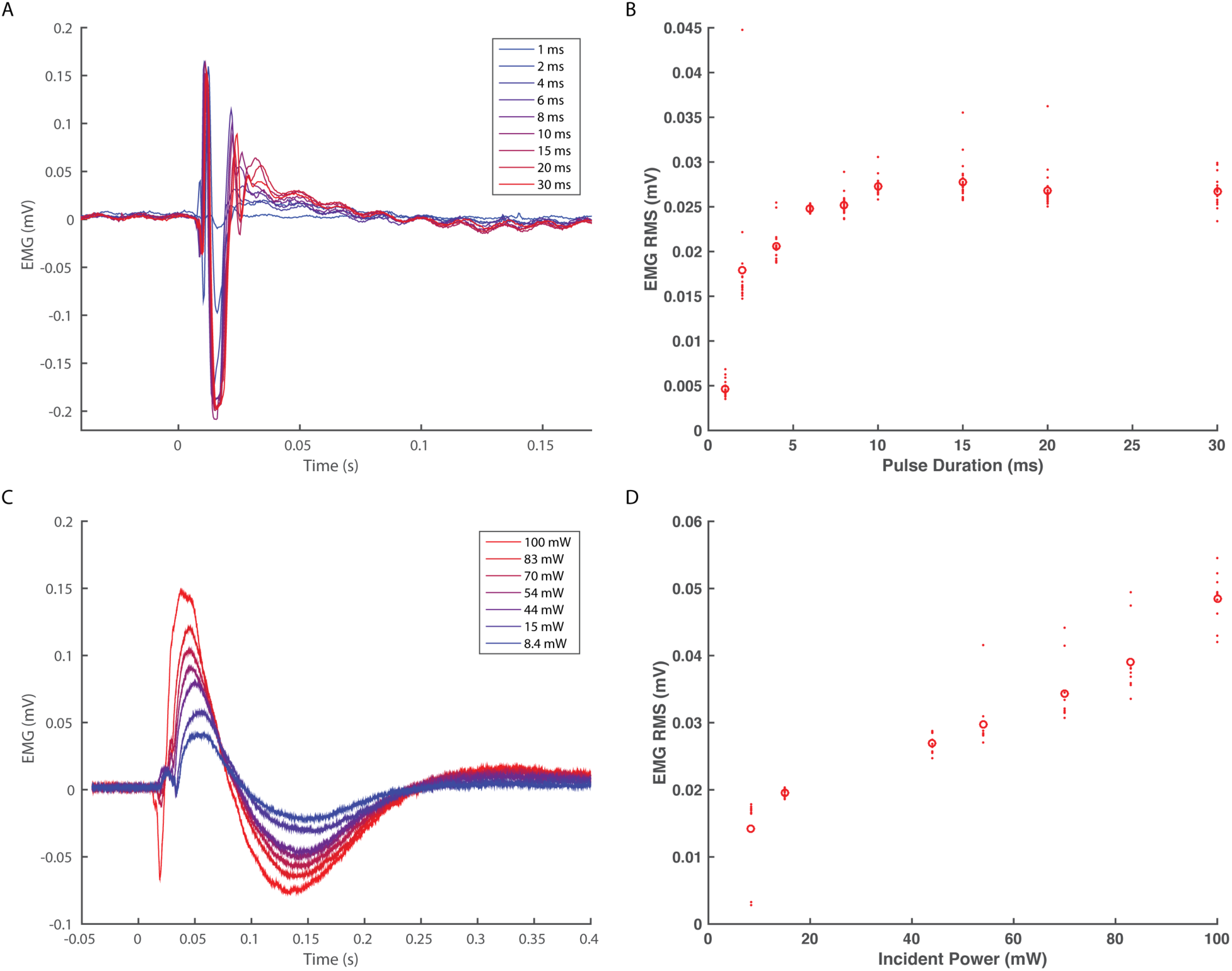
EMG response characteristics to optogenetic stimulation. *A-B) EMG response to varying optical pulse duration*. Panel A shows stimulus averaged EMG traces color-coded by optical pulse duration varying from 1-30 ms (20 pulses, 2.5 Hz trains) while panel B shows the corresponding distributions of RMS values. *C-D) EMG response to varying optical power*. Panel C depicts stimulus-averaged EMG waveforms color-coded by optical power measured at the output of the optical fiber. Stimulus trains consisted of 20 pulses of 20 ms duration at 2.5 Hz. Panel D illustrates the corresponding trend in EMG RMS vs. optical power. Data from A and B are taken from monkey M’s right TA muscle, while data in panels C and D are derived from the left TA muscle. Open circles in B and D indicate mean RMS values while dots indicate RMS responses for individual optical pulses.

As we injected each TA muscle in monkey M with different viral loads approximately an order of magnitude apart (1.66x10^13^ vp in the right TA vs. 2.08x10^12^ vp in the left TA), we examined whether viral load impacted viral transduction and optically elicited muscle activity. We compared the EMG RMS activity in each leg elicited by similar trains (20 ms pulses, 100 mW, 2.5 Hz). Although visual observation did not suggest distinct differences in the magnitude of muscle contractions, the EMG recorded from each leg showed appreciable differences in the shape and duration of the stimulus-averaged waveform. The right TA demonstrated a sharp, transient spike lasting less than 100 ms (Figure 2A) while the left TA demonstrated a waveform lasting 250 ms (Figure 2B). Counterintuitive to the trend expected with respect to viral load, these waveforms correspond to EMG RMS values of 0.027 mV and 0.051 mV, respectively. As we observed above that optical sensitivity was not consistent along a nerve, these differences could arise due to the accessibility of labeled fibers at a given location as opposed to the total number of transduced nerve fibers. In general, however, the range of viral loads injected in this study did not appear to directly correlate with differences in optically stimulated EMG activity.

### 3.4 EMG response to optical pulse trains

After measuring basic EMG responses of optogenetically labeled nerves to single pulses of varying duration and intensity, we then examined the response to longer trains. EMG activity was measured over 10 second blocks of continuous stimulation (20 ms, 100 mW) at increasing pulse frequencies from 2-30 Hz. The train of responses within a frequency block (RMS value of 600 ms window following onset of light pulse) was then normalized to the response of the block’s first stimulus pulse to assess how well the nerve and corresponding muscle activation could track the optical stimulus.

As shown in Figure 3, the EMG response tracked optical stimulation relatively well for pulse frequencies below 16 Hz, retaining EMG responses near 50% of their maximum initial response. Between 16-20 Hz, however, stimulus tracking appears to suffer as the normalized response drops precipitously. At 20 Hz, occasional EMG responses near 50% are interspersed throughout the train from 2-10 seconds, but these are dominated by weak EMG spikes as the nerve/muscle fail to recover. Monkey P demonstrated a similar frequency response with a noticeable dropoff in EMG-optical stimulus coupling between 12-16 Hz (see Supplementary Figure S1c). These results suggest a functional maximum stimulation frequency below 20 Hz, similar to reports of the frequency response of ChR2 in neuronal culture (Boyden et al., 2005; Mattis et al., 2012; Nagel et al., 2003).

**Figure 3.**
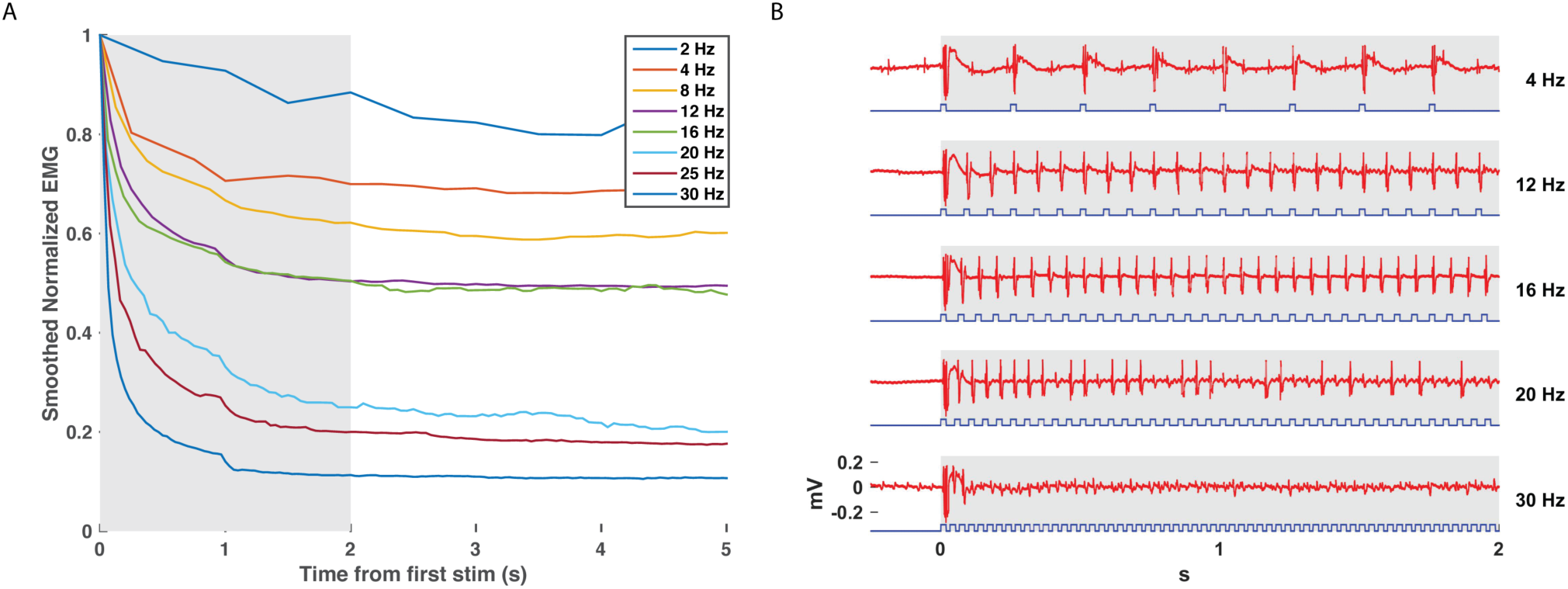
EMG tracking of optical stimulation trains. The right DP nerve of monkey M was stimulated with trains of varying frequency (2-30 Hz, 20 ms pulse width, 100 mW) for 10 seconds while recording EMG from the corresponding TA muscle. The RMS value of the EMG activity elicited by each stimulus pulse (600 ms window following pulse onset) was calculated and normalized by the RMS elicited by the first pulse in the train. Panel A depicts the smoothed EMG response to prolonged optical stimulation at various frequencies, while Panel B depicts the first two seconds of raw EMG responses at selected frequencies from the highlighted window in Panel A. Although a significant drop in elicited EMG activity within the first 1-2 seconds is present at each frequency, stimulus trains above 12 Hz are able to maintain normalized EMG activity at or above 50% of first stimulus magnitude. Between 16-20 Hz, however, elicited EMG waveforms become more erratic as some spikes are missed. EMG responses to stimulus trains above 20 Hz drop off precipitously to below 20% of first stimulus response magnitude.

### 3.5 EMG shows delayed decay with prolonged optical stimulation

Finally, we assessed for any decay in optical sensitivity following optical stimulation in a transduced nerve in Monkey M. The right DP nerve was stimulated continuously via the blue laser at maximum power with a 10 Hz, 20 ms optical pulse train for 2 minutes. Figure 4 depicts the raw EMG trace from monkey M’s right TA muscle as well as the normalized RMS response to stimulation over time. The normalized response falls to 70% of maximum within a few seconds and then levels off similar to the 8 Hz trace in Figure 3. However, after 40 seconds, the muscle response again trends gradually downward over the next 80 seconds before approaching 40% of the initial EMG RMS response at the end of stimulation. Optical stimulation of the right DP nerve of Monkey P showed a similar profile with an initial drop in EMG rms after the initial few pulses followed by a sustained, consistent EMG activity for the rest of the 2 minutes (see Supplementary Figure S1d). The slow decline observed in this study is again consistent with the delayed time course of muscle fatigue with optical stimulation observed in rodent studies (Llewellyn et al., 2010).

**Figure 4.**
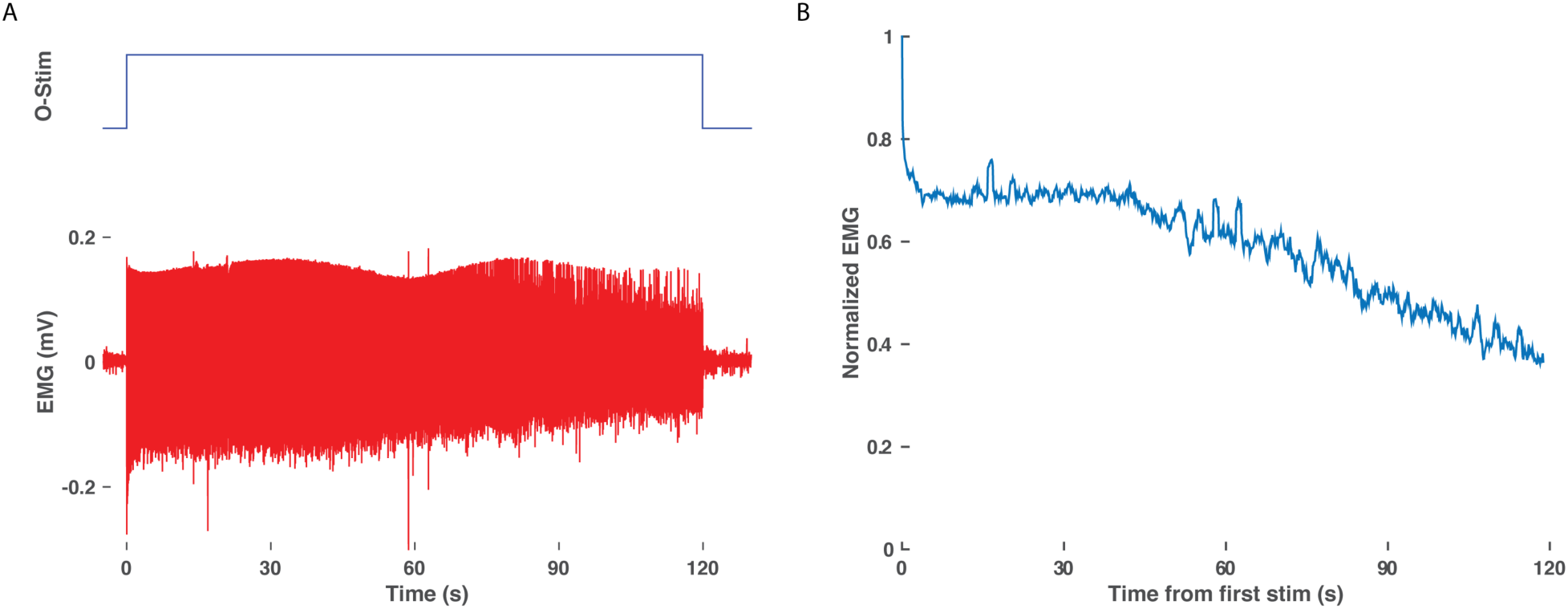
Delay in EMG decay with prolonged optical stimulation. A) Raw EMG traces (bottom, red) from the right TA of monkey M in response to two minutes of optical stimulation (top blue trace, duration of train of 20 ms pulses at 10 Hz). B) Normalized EMG response over time. The RMS response to each optical stimulation pulse was calculated over a 600 ms window following the onset of each pulse and then normalized to the response to the first pulse of the two minute train.

### 3.6 Whole tissue imaging demonstrates variable opsin expression

After final evaluation of functional expression, nerve samples were harvested, cleared, and imaged as whole samples using ribbon confocal microscopy to examine opsin expression patterns. Figure 5a depicts native eYFP fluorescence of an intact whole nerve sample from the right DP nerve of Monkey M, while no similar fluorescence of fiber tracts was observed in a control nerve from an uninjected muscle as seen in Figure 5B. Imaris software was used to trace the eYFP expression in Figure 5A and approximate a longitudinal profile of expression. 3D surfaces corresponding to positive eYFP expression were first computed using a built-in local background signal subtraction algorithm and manual removal of noisy features, with the resulting surfaces highlighted in Figure 5C. The volume of these surfaces was then binned as a function of distance along the length of the nerve (200 μm bins), and the resulting longitudinal profile of opsin/eYFP expression was plotted in Figure 5D. As seen from Figure 5C and 5D, expression of the viral gene product was not uniform along the nerve as patches of expression would emerge and disappear along the nerve. This finding corroborated the previously described variability in the nerve’s sensitivity to optical stimulation (Section 3.1) as well as similar observations in our parallel rodent experiments (see Supplemental Movie 1 (Williams et al., 2016)).

**Figure 5.**
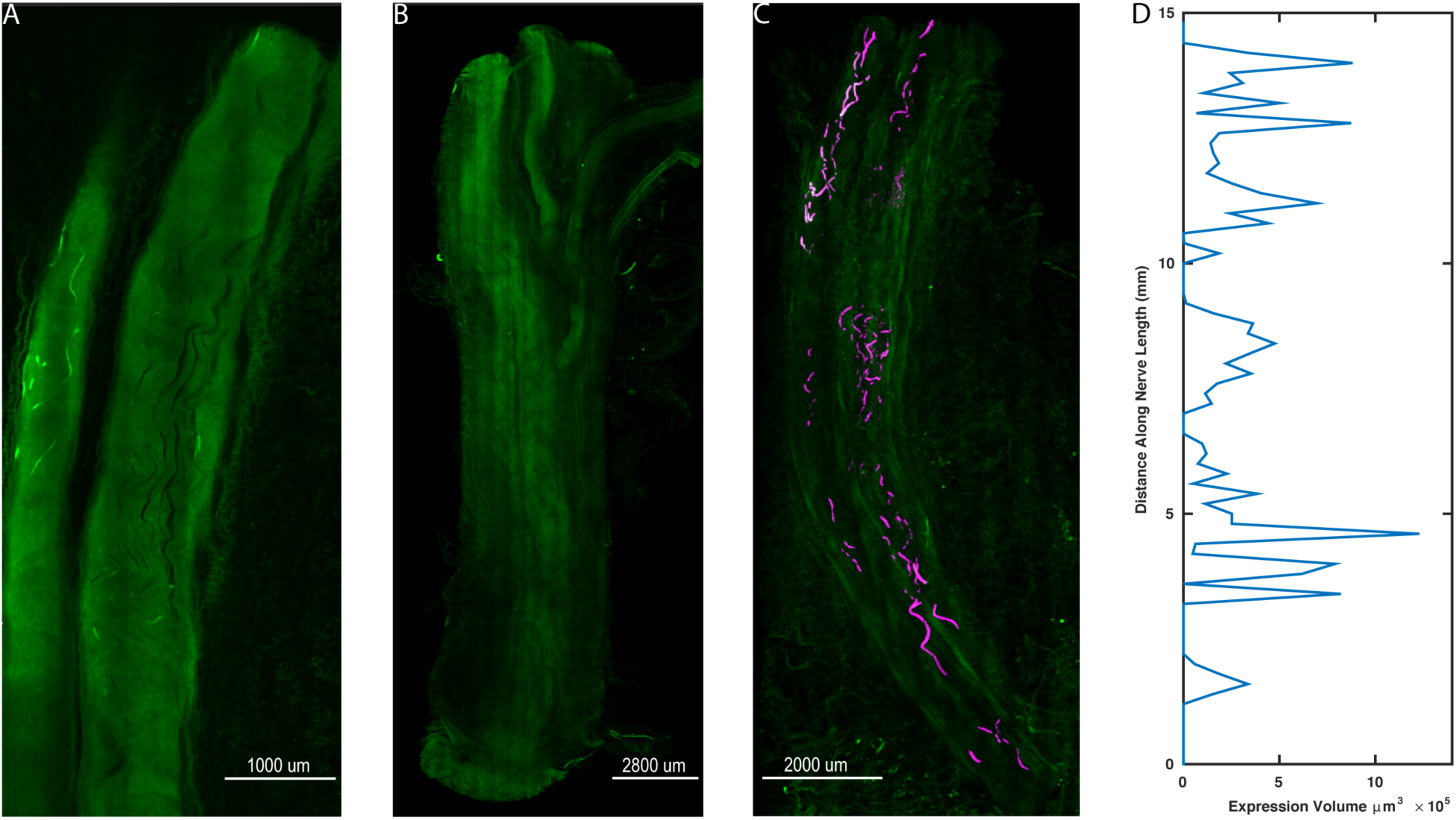
Spatial variability of transgene expression in cleared whole nerve sections. A). Native eYFP fluorescence observed in a cleared nerve section from the right DP nerve of Monkey M using ribbon confocal microscopy. Fluorescent labelling of individual axonal fibers appears to fade in and out within the displayed segment. B) Control nerve from non-injected muscle. In contrast to (A), no distinct patterns of increased fluorescence along individual axons are discernable. C) Maximum intensity projection zoomed-out view of the cleared nerve segment in (A) with expression sites marked. eYFP positive axon segments were labeled in software as 3D surfaces (purple) to highlight the variable labeling of axons along the length of the nerve segment. D) Volume of eYFP expression (horizontal axis) as a function of longitudinal distance (vertical axis) along the nerve in (A) and (C). The dependent variable axis (distance along nerve length) has been rotated to the vertical axis to roughly align with the nerve in (C) and displays how viral expression varies along the length of the nerve.

## 4 Discussion

This study represents a critical step in translating the potential of virally mediated peripheral optogenetics to a clinical therapy capable of alleviating a number of motor diseases or injuries. We have demonstrated that the AAV6-hSyn-ChR2 vector previously shown to be efficacious in transducing peripheral motor axons in rodents following muscle injection (Maimon et al., 2017; Towne et al., 2013) is also a viable vector for peripheral expression of light-sensitive opsins in non-human primates. Our results also exhibit several of the suggested benefits of peripheral optogenetic stimulation over electrical stimulation of muscle activity including graded muscle activation and delayed muscle fatigue. Finally, the correlation of EMG responses to basic optical stimulation parameters lays a foundation from which to approach the design of functional optical stimulation paradigms. Although this study is an important proof-of-concept demonstration, our results also highlight several of the necessary hurdles to be addressed prior to clinical viability as well as new potential avenues of investigation.

### 4.1 Time course of opsin expression

Because we employed a novel longitudinal study of nerve expression in Rhesus monkeys with periodic checks of optical sensitivity, we were able to construct a gross timeline of expression in each animal for comparison with other studies and species. Towne *et al.* utilized a 4-6 week incubation period prior to assessing expression of ChR2 following intramuscular virus injection in rats (Towne et al., 2013). Similarly, a four week incubation period was utilized prior to evaluating the expression of eGFP in the spinal cord following intramuscular injection of AAV6-CMV-eGFP in African green monkeys (Towne et al., 2009). However, a recent study utilizing transdermal stimulation of ChR2-labeled nerves mediated by AAV6 (Maimon et al., 2017) suggests that peak transgene expression may occur later in rats between 5-8 weeks, although even these gross time points of peak sensitivity showed considerable variability. Our findings agree with this variable and potentially extended time course of expression as optical sensitivity was observed initially at 5 ½, 8, and 10 weeks post-injection. In the case of monkey P, although expression was evident relatively early at 5 ½ weeks, optical sensitivity had disappeared by the next check at 9 weeks. As we did not test each injected muscle during earlier evaluations in the first two monkeys in order to minimize surgical manipulations at a given site, we cannot rule out that some sites may have demonstrated optical sensitivity at earlier time points similar to Monkey P. Additionally, the focal sensitivity observed along the left DP nerve of monkey M raises the possibility we did not fully expose or probe one of these focal “hotspots” of sensitivity during our earlier assessments while attempting to leave the surrounding tissue grossly intact. Once we more aggressively exposed the DP nerve and its insertion into the TA muscle, stimulation of one of these hotspots likely became more probable. In any case, the time course of expression, as well as differences between species, remains a critical yet poorly understood process.

### 4.2 Considerations for chronic optical stimulation

Our results also bring forth several considerations for chronic FOS. As one potential application of this gene therapy is to restore volitional control of paralyzed muscle activity through a hybrid optogenetic-BMI, optical nerve stimulation hardware such as chronic LED or fiber optic nerve cuffs must be able to consistently stimulate opsin-labelled axons over a period of years. A potential benefit of using chronically implanted optical nerve cuffs on virally targeted nerves would be the ability to assess the time course of expression without additional surgical procedures. However, the variable expression and sensitivity pattern of ChR2 observed in monkey M in Figure 5 and some of our parallel rat studies (see Supplemental Movie 1 (Williams et al., 2016)) suggests that proper placement of stimulation hardware for either of these applications may be more challenging than initially anticipated. Correct temporal assessment of opsin expression patterns would require blind, accurate placement of nerve cuffs soon after injection over high expression zones on the nerve. Similarly, to provide consistent chronic optical stimulation capabilities in a rehabilitation setting would require 1) an additional evaluation surgery following the virus incubation period to properly place optical cuffs, 2) securing the cuff such that it does not move relative to the hotspot of expression on the nerve, and 3) stability of expression/low turnover at the hotspot. It is possible that the variable optical sensitivity and fluorescent expression observed in this study is due in part to poor expression and trafficking of opsins to the axonal membrane, although another likely contributor may be the immune system through a piecemeal recognition and degradation of opsins by an immune response. Elucidating the underlying cause of this problem could then direct further development of opsins or promoters (Chaffiol et al., 2017) with better expression and trafficking characteristics versus development of injection techniques (Burger et al., 2005; Favre et al., 2000; Harris et al., 2012; Tosolini and Morris, 2016; Williams et al., 2016), recombinant viruses (Bartel, 2011; Tervo et al., 2016) to increase the efficiency and total number of axons transduced within a nerve, or immunosuppressive approaches to maintain expressed opsins.

### 4.3 Transduction as a function of viral load

As we contemplate potential approaches to improve opsin expression along the nerve, we must also examine the results of this study with respect to viral load delivered to the muscle. We used a range of viral loads spanning approximately an order of magnitude (∼10^12^-10^13^ vp per muscle) between the three monkeys with no obvious trend in expression. The lower end of this range is consistent with a previous study by Towne *et al* in African green monkeys (Towne et al., 2009). However, the range of viral loads per kilogram of bodyweight used in this study (9.24x10^11^-1.84x10^12^ vp/kg) is an order of magnitude lower than the high titer intramuscular injections used by Maimon et al. in rats (1.5-2.5x10^13^ vp/kg) (Maimon et al., 2017). Assuming that the mass and volume of the muscles targeted in these studies scale approximately with total body weight, a lack of functional limb movement from optical stimulation in this study could be explained by an insufficient dose of viral particles delivered to muscles with much greater volume compared to prior mouse and rat models. Based on rodent studies, it is possible that doses on the order of 10^14^ vp per muscle might be necessary to yield consistent opsin expression that is functional for eliciting limb movements in primates.

### 4.4 Virus delivery approaches

The differences in viral load as a function body weight across animal models highlights another difficulty in scaling this gene therapy approach up to humans. As the volume of muscle and corresponding zone of neuromuscular junctions targeted for viral uptake increases dramatically from rodent to primate, efficient delivery of viral particles to the entire motor end plate may become both expensive and technically challenging. Our first attempt to address this challenge was to simply increase the volume of viral solution injected with hypertonic saline to be on the same order of magnitude used in rodent studies (∼200 μL/kg bodyweight vs 100 μL/kg bodyweight in (Maimon et al., 2017)) with the potential ramifications discussed above. Our second approach was to attempt to localize zones of high neuromuscular junction density near the motor end plate using electrical stimulation. Previous rodent studies have demonstrated that targeting muscle injections along motor endplates greatly enhances motor neuron transduction (Tosolini et al., 2013; Tosolini and Morris, 2016). Targeting of the motor end plate as in these studies requires prior histological mapping of the motor end plate in a given muscle *in situ* followed by visual alignment of anatomical landmarks in the subject to be injected. Conversely, our approach uses electrophysiological responses to map the end plate and potentially account for anatomical variability between animals. A third injection approach that we employed in Monkey P that may be promising for scaling up injections with animal size was to inject virus directly into the nerve branch of interest. Our experience with this technique has shown that intraneural injections near the insertion of the nerve into the muscle may effectively utilize the nerve sheath to contain and funnel the virus toward the motor end plate as the nerve branches out within the muscle (Williams et al., 2016). Therefore, virus that does not directly enter nerve axons upon injection but instead resides in the connective tissue perineurium may still be guided back down to the muscle where it may have a greater probability of uptake at neuromuscular junctions. Utilization of nerve injections in this manner could significantly reduce the volume of virus needed for effective motor neuron transduction in larger animals such as the macaque. Intraneural injections at sites more proximal to the spinal cord such as the sciatic nerve do hold the possibility of transducing unwanted sensory neurons. However, injecting the nerve closer to the target muscle would likely minimize unrelated sensory transduction or limit it to proprioceptive fibers that could be utilized for feedback.

### 4.5 Viral vector design

In addition to the load and route of viral particles delivered to motor nerves, the composition of the viral construct itself holds great potential for improvement of motor nerve transduction. The mechanisms by which AAV vectors undergo uptake at the neuromuscular junction and traffic to the spinal cord are not completely understood, but presumably it is a receptor-mediated process facilitated by domains on the viral capsid which confer tissue tropism to various serotypes. A better fundamental understanding of these uptake and transport processes could inform the design of viral vectors for peripheral motor gene therapies. An alternative approach recently taken by several groups is “directed evolution” or high-throughput screening and selection of recombinant AAV variants for a desired trait (Choudhury et al., 2016; Dalkara et al., 2013; Tervo et al., 2016). Similarly, the hSyn promoter has been commonly used for peripheral nerve transduction due to its specificity for neural tissues yet relatively strong expression. Using a promoter restricting expression to specific nerve fiber types (e.g. slow/fast fatiguable motor units, proprioceptive fibers, etc.) would enable selective modulation of efferent or afferent activity as well as an approach to artificially specify the recruitment order of motor unit types. However, in general, more specific promoters result in weaker expression in the target tissues, so this tradeoff of nerve optical sensitivity versus fiber type specificity would have to be addressed.

### 4.6 Immune Response to AAV

AAV6 was chosen as the gene delivery vehicle in this study due to AAV’s safety profile and low immunogenicity (Calcedo and Wilson, 2013) as well as its previously demonstrated success in transducing peripheral motor nerves in non-human primates (Towne et al., 2009). However, a considerable proportion of both humans and macaques naturally exhibit pre-existing neutralizing antibodies (NAbs) to a variety of AAV serotypes (Boutin et al., 2010; Calcedo and Wilson, 2013; Hurlbut et al., 2010). Even at low NAb levels, transgene expression may be significantly inhibited in non-human primates (Hurlbut et al., 2010; Jiang et al., 2006). We did not assess the status of preexisting NAbs to AAV6 in our subjects, so its role regarding differences in expression between monkeys in this study is unclear. Nevertheless, several transient immunosuppression strategies such as those used for organ transplants have shown efficacy in maintaining AAV6-mediated transgene expression in canine models (Shin et al., 2012; Wang et al., 2012) as well as non-human primate models examining AAV8 (Jiang et al., 2006). Employing such regimens may not only increase transduction efficiency and prolong transgene expression, it could also enable separate viral injections of multiple muscle groups over several surgeries without decreased efficacy after an initial viral exposure (Riviere et al., 2006).

### 4.7 Optical stimulation parameters and opsin selection

The results from this study provide baseline practical guidelines for optical stimulation parameters. EMG responses were modulated with pulse widths up to 10 ms, above which responses appeared to plateau. From a frequency response perspective, EMG responses tracked optical stimulation trains up 16 Hz, suggesting an upper bound for use in FOS stimulation schemes. This limit is relatively low compared to the frequency of stimulus trains commonly used for FES, often ranging from 20-50 Hz for clinical applications (Doucet et al., 2012). However, due to the previously discussed differences in recruitment order between optical and electrical stimulation, further study is required to elucidate how these optical stimulation parameters translate to functional force production and how to optimize modulation strategies for neuroprosthetic driven movements.

The EMG relation to optical stimulation parameters explored here was only characterized for the opsin ChR2. Although its use in this and similar prior studies in rodents as a first line of investigation is warranted by ChR2’s well-characterized behavior and consistent expression patterns in a wide array of neural systems, other recently developed opsins may hold properties beneficial to peripheral motor stimulation. Indeed, we injected several muscles in our second monkey with a construct using the opsin Chronos to exploit Chronos’s increased sensitivity and faster kinetics to 1) lower the light intensity and consequently power requirements for implantable optical stimulation hardware, and 2) increase the frequency range of pulsed stimulation trains to at least comparable levels used for FES. Recent studies have supported the fast temporal advantages of Chronos over ChR2 in the central auditory pathway (Guo et al., 2015; Hight et al., 2015). Although this study demonstrated a novel use of Chronos in the motor periphery, the brief period during which we were able to observe its response left us unable to fully examine whether these purported benefits extend to the peripheral motor system. However, preliminary data from our parallel rat studies (Williams et al., 2016) suggest that Chronos does have a better frequency response for light stimulus-EMG coupling in the periphery than ChR2 (see Supplemental Figure S2). Other opsins which may prove beneficial for peripheral applications include red-shifted variants such as Chrimson (Klapoetke et al., 2014). The use of longer stimulation wavelengths would allow greater tissue penetration that could prove highly desirable when scaling stimulation hardware up to target primate nerves several millimeters in diameter.

### 4.8 Comparison with spinal electrical stimulation approaches

Our main goal in this study was to introduce peripheral optogenetic stimulation as an alternative to FES in BMI applications. FES is typically associated with electrical stimulation of muscles directly or through nerve stimulation. However, several groups have used brain-controlled electrical intraspinal (Zimmermann and Jackson, 2014) or epidural spinal (Capogrosso et al., 2016) stimulation in NHPs for grasping and hindlimb locomotion, respectively. This mode of stimulation elicits muscle activation patterns either directly by stimulating pools of alpha motoneurons or indirectly by inducing motor patterns through interneurons following stimulation of dorsal roots. Thus, our peripheral FOS approach may not be directly comparable to electrical stimulation approaches at the spinal level. Nonetheless, it is possible that the potential benefits of FOS observed in the periphery may also extend to analogous optical stimulation of the spinal cord as has been recently investigated (Mondello et al., 2018).

## 5 Conclusions

In summary, the viral transduction and functional expression of opsins for peripheral optical modulation of muscle activity in non-human primates is a step toward efficient reanimation of movement in paralyzed subjects. The introduction of neuromuscular junction targeting for virus injection is a useful technique for increasing the likelihood of virus uptake. In addition, the EMG response characteristics to optical stimulation parameters described here serve as an important base upon which to build future primate studies and FOS algorithms.

While the jump from rodent to primate is important in itself, this study also highlights problems due to differences in scale and species that may not have been as pronounced in prior rodent studies. Potential variability in both the timeline and spatial profile of expression, the immune system’s probable role in this variability, and effectiveness of the virus as well as light delivery in much larger target muscles/nerves are all challenges that must be addressed before FOS may become a clinically viable approach to restoring lost motor function.

## Supporting information

## 8 Acknowledgements

We would like to thank Chris Towne for his consultation regarding peripheral optogenetic techniques. We thank Richard Dum, Jean-Alban Rathelot, and Peter Strick for their guidance and expertise with primate virus injections, Doug Weber for his aid with EMG recordings, and Bistra Iordanova for her help with histology. Finally, we would like to thank Kathy Hansell-Prigg, Scott Kennedy, Hongwei Mao, Steve Suway, Rex Tien, and Sally Zheng for their assistance with surgeries and experiments.

Note: A version of this manuscript was previously released as a pre-print (Williams et al., 2018) to the bioRxiv server hosted by the Cold Spring Harbor Laboratory.

## 9 Author Contributions

JW, AV, and AS contributed to study conception and experimental design. JW performed all surgeries and experiments. AW performed tissue clearing and imaging while JW and AW analyzed imaging data. JW wrote the first draft of the manuscript. All authors contributed to manuscript revision, read and approved the submitted version.

