## Supplementary material for "Viral-mediated optogenetic stimulation of peripheral motor nerves in non-human primates"

### 1 Supplementary Methods

In a set of experiments parallel to our non-human primate experiments, male Fisher 344 rats (200-400 grams, Charles River Laboratories, Wilmington, MA) were injected with the same AAV constructs used in the monkey study to verify construct expression and EMG response characteristics as well as to refine injection techniques in muscle, nerve, or spinal cord.

#### 1.1 Target Muscles

The tibialis anterior (TA) muscle of one leg was targeted for each rat. The fascia was typically cut if visualization of the corresponding nerve was necessary or if injecting into the nerve itself, and the fascia was sutured following injection. Otherwise, muscle injections were performed through the overlying fascia. Spinal cord injections targeted the spinal cord level of the TA muscle (L4-L5) by injecting through a burr hole at the T13-L1 vertebral level of the spine. The anterior horn of the spinal cord was targeted using low-level electrical stimulation through a 30 gauge monopolar injectable needle while monitoring TA and surrounding muscle contractions.

#### 1.2 Virus Injections

Two high-titer AAV6-based viral vectors were obtained from Virovek, Inc. (Hayward, CA) and used in rat injections. Either an AAV6-hSyn-ChR2(H134R)-eYFP construct ($1.04 \times 10^{14}$ vp/mL) or AAV6-hSyn-Chronos-eYFP ($1.00 \times 10^{14}$ vp/mL) were injected at a given target site. Virus was injected either undiluted or diluted to twice its normal volume with hypertonic saline. Injections were performed using a Hamilton syringe and a programmable syringe pump. A 30 gauge needle was used for muscle and spinal cord injections, while a 35 gauge needle was used for nerve injections. Muscles were injected with 15-30 μL of virus at 5 μL/min, nerves with 3-5 μL of virus at 2 μL/min, or the spinal cord with 2-3 μL of virus at 0.5 μL/min. SR101 (sulforhodamine) was occassionally mixed with the viral solution to confirm successful injection.

#### 1.3 Functional evaluation of opsin expression

Peripheral AAV injections were allowed to incubate for 3-6 weeks following injection prior to evaluating construct expression. Spinal cord injections were allowed to incubate for 1-2 weeks prior to evaluation. During an evaluation surgery, optical stimulation was provided by a 473 nm laser through an optical fiber held on a micropositioner. To record EMG responses, a pair of 36 AWG stainless steel wires, stripped and barbed at the end, was inserted into the muscle belly of the target muscle. A common ground wire was inserted into the base of the rat's tail. EMG signals were sampled and recorded at 25 kHz using a Tucker Davis Technology Systems neurophysiology base station (RZ5D, TDT, Alachua, FL).

### 2 Supplementary Data:

**<u>Supplementary Movie 1. Variable optical sensitivity along virally-transduced rat nerve.</u>** The movie clip demonstrates variable sensitivity to optical stimulation along the length of a rat nerve expressing ChR2 following muscle injection of AAV6-hSyn-ChR2-eYFP. The 472 nm wavelength optical beam is initially centered over the injected tibialis anterior muscle, showing no muscle response to optical stimulation. As the beam path begins to travel along the transduced nerve, small muscle contractions corresponding to optical stimulation slowly grow in magnitude as the beam approaches an expression "hot spot". Toe and ankle movements are also gradually added. The contractions grow to a maximum over the hot spot and then die away as the beam travels away. This pattern reverses as the beam retraces its path back toward the muscle.

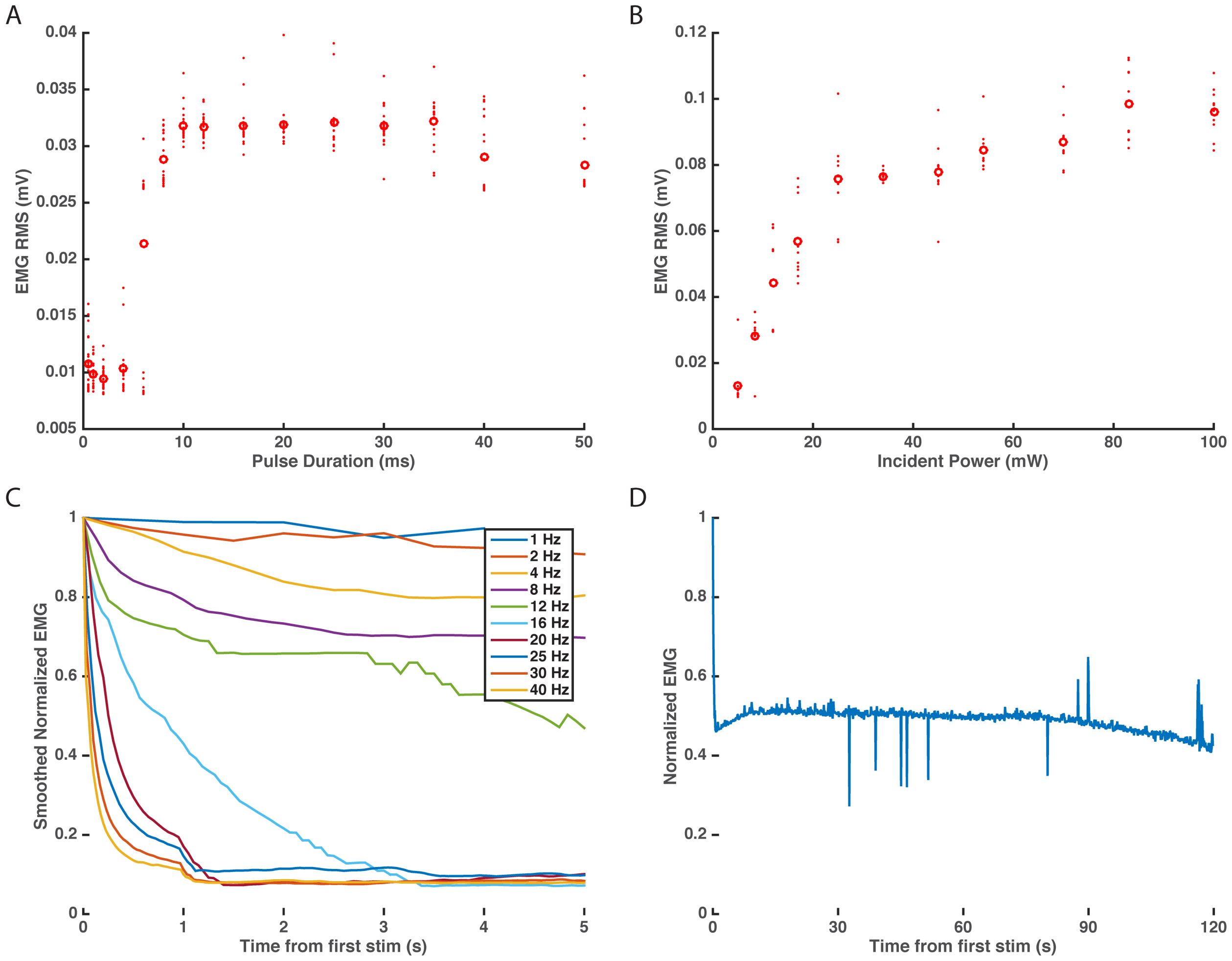


**Supplementary Figure S1. Optical stimulation results from Monkey P.** A) EMG vs. optical pulsewidth. B) EMG vs. incident power. C) Frequency response of EMG-optical stimulus coupling. D) EMG response to prolonged, continuous optical stimulation (20 ms pulsewidth, 10 Hz, 2 minute trains x 2). Results agree with corresponding trends from Monkey M displayed in Figures 2-4 of the main text.

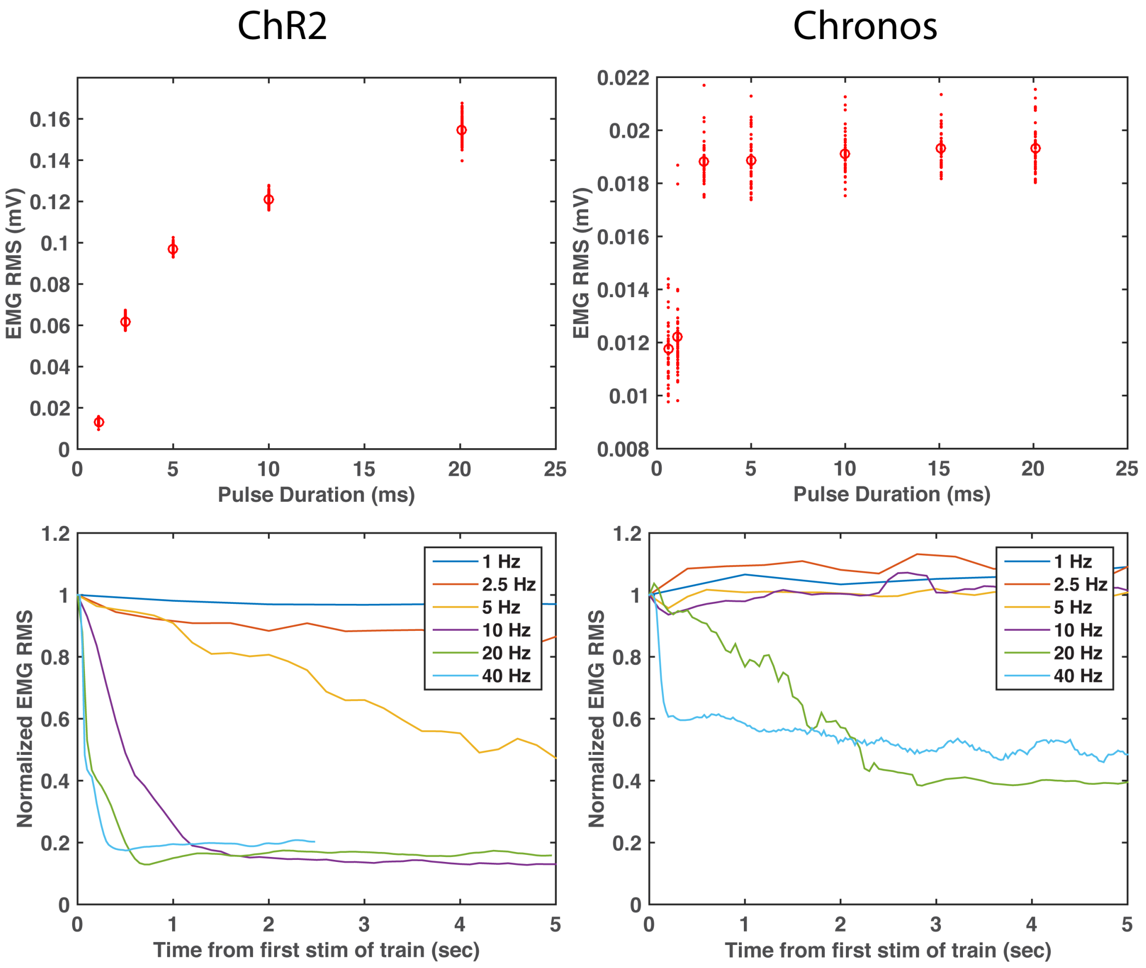


**Supplementary Figure S2. Comparison of ChR2 vs. Chronos in transfected rat motor neurons for muscle stimulation.** Top row: EMG response from ChR2 (left) and Chronos (right) stimulation as a function of optical pulse duration. Chronos shows increased sensitivity over ChR2 at lower light levels (shorter pulses). Bottom row: EMG tracking of optical pulse train. Colored curves show the average EMG response (normalized to the first optical pulse response) to a train of 10 ms pulses over time at indicated frequencies. ChR2 EMG responses begin to lose the ability to track optical stimulation above 20 Hz, while Chronos responses still maintain some tracking to prolonged stimulus trains at 40 Hz.
